# Engineering a minimal SilkCatcher/Tag pair compatible with SpyCatcher/Tag pair for the production of native-sized spider silk

**DOI:** 10.1101/2022.10.21.513168

**Authors:** Ruxia Fan, Johanna Hakanpää, Karoliina Elfving, Helena Taberman, Markus B. Linder, A. Sesilja Aranko

## Abstract

Protein/peptide pairs, called Catcher/Tag pairs, are applied for biological isopeptide-bond mediated Click-reactions. Covalent protein ligation using Catcher/Tag pairs has turned out to be a valuable tool in biotechnology and biomedicines. It is essential to increase the current toolbox of Catcher/Tag pairs to expand the range of applications further, e.g., for multiple-fragment ligation that requires several orthogonal ligases. We report here engineering of novel Catcher/Tag pairs for protein ligation, aided by a new crystal structure of a minimal CnaB domain from *Lactobacillus plantarum*. We engineer several split variants, characterize in detail one of them, named SilkCatcher/Tag pair, and show that the newly engineered SilkCatcher/Tag pair is orthogonal and compatible with the widely used SpyCatcher/Tag pair. Finally, we demonstrate the use of the new SilkCatcher/Tag pair in the production of native-sized highly repetitive spider-silk-like proteins with >90% purity, which is not possible with the traditional recombinant production.

## Introduction

Advances in the methods of covalent ligation of proteins *in vivo* and *in vitro* have expanded the possibilities of recombinant production and created numerous applications, such as engineering non-linear architectures, as well as protein semisynthesis, cyclization and purification ^[1–5]^. Protein ligation using Catcher/Tag pairs has obtained wide attention since the first reports in 2010 due to the efficiency and robustness of the reaction ^[6–8]^. The method is based on autocatalytic formation of an isopeptide bond between a Cather/Tag protein/peptide pair complex. The Cather/Tag pairs are derived from CnaB domains, Ig-like domains found in collagen binding cell surface proteins in gram-positive bacteria, which carry intramolecular isopeptide bonds that stabilize the structure ^[9,10]^. Some CnaB domains can be split into two fragments; a ∼10 kDa larger fragment, named Catcher, and a ∼1.5-2.5 kDa peptide, called Tag, that associate together by protein fragment reconstitution. The association is followed by an autocatalytic isopeptide bond formation, thus leading to covalent conjugation of the protein-peptide pair and proteins fused to them. Elegant engineering combined with directed evolution has further improved the SpyCatcher/Tag pair turning it into a powerful tool for the covalent conjugation of proteins which has been adapted into multiple applications ^[4,11–13]^. The successful engineering of SpyCatcher/Tag pair has inspired search for other Catcher/Tag pairs and several new pairs have been reported ^[14–17]^.

A widely interesting application of protein conjugation is multiple fragment ligation ^[15,18,19]^ that allows the recombinant production of long and repetitive proteins that are difficult to produce in the common production hosts, such as *Escherichia coli*. Spider silk is an important and interesting example of this, because of its extreme properties both at molecular and macroscopic level, as well as the numerous applications ^[20–22]^. The desirable mechanical properties of spider-silk fibers, together with the unfeasibility of farming spiders, have prompted research aiming at producing recombinant spider silk. Spider silk proteins, called spidroins, consist of very long (thousands of amino acids, 250-320 kDa) and highly repetitive alanine- and glycine-rich regions ^[23,24]^. The size and repetitive nature of the core parts are crucial for the fiber properties but create significant challenges for the recombinant production of the spidroins ^[21,22]^.

Multiple fragment ligation is, however, limited by the lack of efficient and orthogonal (non-cross-reactive) protein ligation methods. The use of non-orthogonal methods would likely lead to predominantly intramolecular cyclization reaction ^[25–27]^. In addition, the length of the products of intermolecular conjugation could not be controlled but instead multimers of varying lengths would be formed. The uncontrolled formation of side-products makes the system unsuited for any larger scale production. Thus, a toolbox of orthogonal Catcher/Tag pairs is needed. In addition to being orthogonal, these pairs should preferably work in the same conditions, and be robust in terms of the target proteins.

In this work, we report the engineering of novel Catcher/Tag pairs starting from two CnaB domains from a probiotic bacterium, *Lactobacillus plantarum*. We solved the crystal structure of one of the CnaB domains and discuss the design principles for split proteins. We optimize the reaction conditions for the most promising pair, that we here named SilkCatcher/Tag. We demonstrate that it can be used for a pH switch, and show that the newly designed pair is orthogonal in binding and therefore forms a compatible pair with the SpyCatcher/Tag pair. Furthermore, we show multiple fragment ligation to produce native-sized spider silk proteins *in vitro* taking advantage of the orthogonality of SpyCatcher/Tag and our newly designed SilkCatcher/Tag.

## Results and Discussion

### Engineering novel Catcher/Tag pairs

A cell surface protein lp2578 from a lactic acid bacterium *Lactobacillus plantarum* was predicted to contain a collagen binding domain and two CnaB domains ^[28]^ (Figure 1A, Figure S1A). Both CnaB domains had residues suitable for isopeptide bond formation suggesting they would contain intramolecular isopeptide bonds (Figure 1B, Figure S1A). We produced the two CnaB domains, named CnaB1 and CnaB2, recombinantly in *Escherichia coli* as fusion proteins with N-terminally H6-tagged Smt3 -tag, which was subsequently cleaved of with Ulp1 (Figure S1B). Both CnaB domains were expressed in high yields and were soluble at high concentrations. The Far-UV CD spectra of the two proteins were characteristic to CnaB domains with a positive peak at ∼232 nm and a negative peak at ∼216 nm (Figure S1C). Furthermore, the molecular weights of the CnaB1 and CnaB2 domains obtained by mass spectrometry corresponded to the sizes expected for a product containing one isopeptide bond each (Figure S1D, E).

**Figure 1.**
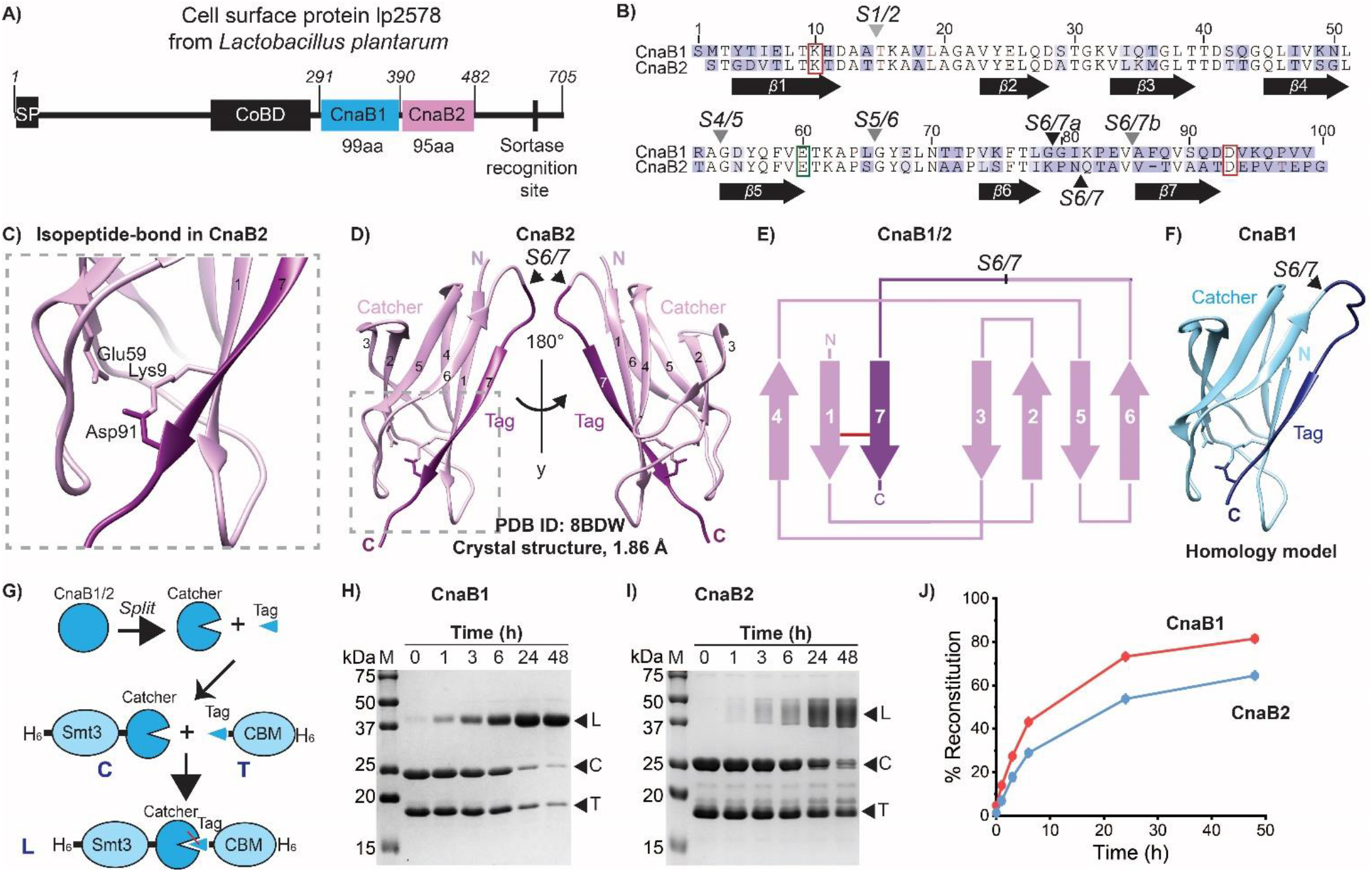
Engineering Catcher/Tag pairs from the *L. plantarum* CnaB1 and CnaB2 domains. A) Schematic presentation of the domain structure of *L. plantarum* cell surface protein lp_2578. Key amino-acid sequence-numbers are shown. SP stands for signal peptide, CoBD stands for collagen binding domain. B) Sequence alignment of CnaB1 and CnaB2 domains. Lys and Asp participating in the reaction are highlighted with red boxes and the catalytic Glu with a green box. Split sites (S) are numbered according to the flanking β-sheets (preceding/following) and indicated with arrowheads. C) Isopeptide bond in the newly solved *L. plantarum* CnaB2 structure (PDB ID: 8BDW). D) Cartoon presentation of the crystal structure of *L. plantarum* CnaB2 domain. Tag (dark purple) and Catcher (light purple) regions of the first tested split variants are shown. β-sheet numbers are given and the side chains of the active site residues are shown with stick model. Split sites are shown with arrow heads. E) Topology diagram of *L. plantarum* CnaB1 and 2 domains based on the homology model and crystal structure. Tag (dark purple) and Catcher (light purple) regions of the first tested split variants are shown. F) Cartoon presentation of the homology model of *L. plantarum* CnaB1 domain. Tag (dark blue) and Catcher (light blue) regions of the first tested split variants are shown. The side chains of the active site residues are shown with a stick model. G) Schematic presentation of engineering split Catcher/Tag pairs from CnaB domains and their fusion to H6-Smt3 and CBM-H6 tags for purification and ligation studies. H and I) Analysis of the time-scale of the ligation of CnaB1(S6/7a) and CnaB2(S6/7) Catcher/Tag pairs on SDS-PAGE. M stands for molecular weight marker. L, C, and T stand for ligation product (H6-Smt3-Catcher-Tag-CBM-H6), Catcher-precursor (H6-Smt3-Catcher) and Tag-precursor (Tag-CBM-H6), respectively. J) Reaction rate of the CnaB1 and CnaB2 S6/7 Catcher/Tag pairs. Molecular graphics were prepared with UCSF Chimera [29].

While mass spectrometry and sequence homology strongly indicated that the two proteins would contain isopeptide bonds, we wanted to obtain more detailed structural information to aid the design of a Catcher/Tag pair. We solved the crystal structure of the CnaB2 domain with 1.86 Å resolution (Figure 1C, D, Supplementary Table 1). CnaB2 domain had an unusual crystal packing forming a hollow sphere, each ball consisting of 8 trimers and altogether 24 domains, with 12 domains within the symmetric unit (data not shown). The CnaB2 domain had typical CnaB-fold consisting of seven β-strands (Figure 1C-F).

As anticipated based on the sequence homology and mass spectrometry data, the CnaB2 domain contained one isopeptide bond connecting the residue Lys9 in the first β-sheet and Asp91 in the last β-sheet (Figure 1C), which was clearly visible in the electron density map. Comparison to the prediction made with AlphaFold ^[30,31]^ shows that although the prediction by AlphaFold is very good, it is missing the isopeptide bond. We were not able to obtain diffracting crystals from the CnaB1 domain, most likely due to the extremely high solubility of the protein. Instead, a homology model was built with SWISS-MODEL ^[32]^ based on the structure of the CnaB2 domain, followed by manual adjustment (Figure 1E). Interestingly, CnaB1 (99aa) and CnaB2 (95aa) domains are smaller than the CnaB domains used as starting point for the engineering of the three versions of SpyCatcher/Tag (SpyCatcher/Tag1-3) (121aa), SnoopCatcher/Tag, and DogCatcher/Tag pairs (127aa) (Figure S2).

We engineered Catcher/Tag pairs from CnaB1 and CnaB2 domains by splitting at the loop between the β-strands numbered six and seven (Figure 1D-F). There are no clear guidelines for selecting the split site for protein fragment complementation, and the optimal solution is application-dependent ^[33–35]^. The split site was thus selected based on the design of SpyCatcher/Tag system ^[6,7]^. The Catcher/Tag pairs are called CnaB1Catcher/Tag(S6/7a) and CnaB2Catcher/Tag(S6/7) with the number in parenthesis indicating the location of the split site (Figure 1B). The Catcher/Tag pairs were tested in a model system, in which Catcher was fused to an N-terminal H6-Smt3 tag (H6-Smt3-Catcher), whereas Tag was fused to a C-terminal cellulose binding module (CBM) and a H6-tag (Tag-CBM-H6) (Figure 1F). Both CnaB1 and CnaB2 Catcher/Tag pairs were able to catalyze protein ligation in optimized conditions as observed by analysis with SDS-PAGE, in which a protein corresponding to the size of a ligation product appeared upon mixing (Figure 1G-J). Instead, no ligation was observed when the catalytic, isopeptide-forming Asp in CnaB2Tag was mutated to Ala (FigureS3A). The ligation yield of the CnaB1Catcher/Tag(S6/7a) pair was better than that of the CnaB2 Catcher/Tag pair (Figure 1J), which prompted us to study the newly engineered CnaB1 Catcher/Tag pair in more detail.

Analysis of the ligation product of CnaB1Catcher/Tag(S6/7a) pair by mass spectrometry indicated that an isopeptide bond was connecting the Catcher/Tag pair (Figure 2A).

**Figure 2.**
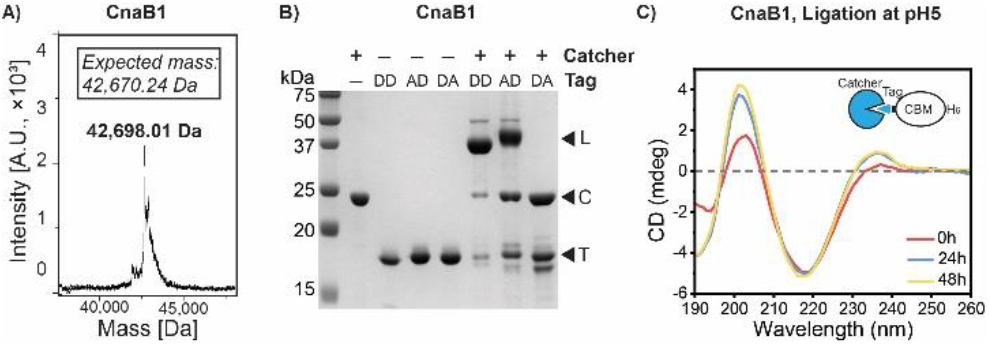
CnaB1Catcher/Tag(S6/7a) ligation. A) Molar mass of the ligation product analysed by mass spectrometry. B) SDS-PAGE gel showing the ligation reaction of the wildtype Catcher with the wildtype Tag (Asp92&Asp93; DD), and the two alanine mutants; Asp92Ala (AD) and Asp93Ala (DA) Tag. L, C, and T, stand for ligation product, Catcher, and Tag. C) CD spectra from ligation reaction of CnaB1Catcher/Tag pair after 0h, 24h, and 48h.

To confirm which one of the two Asp residues located near the predicted isopeptide bond (Asp 92 and 93) was participating in the reaction, we tested ligation reaction after mutating them into alanine one-by-one (Figure 1B, Figure 2B). The mutant harboring Asp93Ala mutation was inactive, whereas the Asp92Ala mutant was active, showing that Asp93 was taking part in the isopeptide bond formation. Finally, formation of the CnaB1 fold during the reaction of CnaB1Catcher/Tag pair could be followed by CD-spectroscopy, in which the characteristic negative peak around 216 nm and positive peak around 232 nm appeared after incubating the Catcher/Tag pair together (Figure 2C).

### Optimization of ligation reaction

We next studied the optimal reaction conditions. Reaction of Tag-precursor could be driven in close to 100% when mixed with excess of the Catcher part (Figure 3A). Reaction was found to work in all tested temperatures 4-45°C, while being optimal at 37°C (Figure 3B). Unlike the CnaB2Catcher/Tag pair (Figure S3B-D and data not shown), the CnaB1 Catcher/Tag pair was robust in terms of the fusion proteins (data not shown) and highly soluble. The reaction had a pH optimum at pH 5, in which ∼80% of the precursors had reacted, whereas at pH 3 and pH 7 the yield was just 30% and at pH 8 and pH 9 only very faint bands for the ligation product were seen (Figure 3C). The ligation reaction of the SpyCatcher/Tag variants 1-3 and another Catcher/Tag pair, called SdyCatcher/Tag^[16]^, are also optimal at pH 5-6 ^[6,11,16]^. All these pairs have Lys and Asp as the isopeptide bond forming residues. In contrast, SnoopCatcher and DogCatcher, in which isopeptide bond is forming between Lys and Asn, have optimal reaction at pH >9 ^[14,15]^. The similar optimal pH to that of SpyCatcher/Tag1-3 provides possibilities for their use in single-pot reactions. Inspired by the drastic differences in activity depending on pH, we wanted to test the possibility to activate the reaction by a pH switch. Indeed, almost no reaction took place after 24h incubation at pH 9, but the activity was retained after lowering pH to 5 (Figure 3D, E).

**Figure 3.**
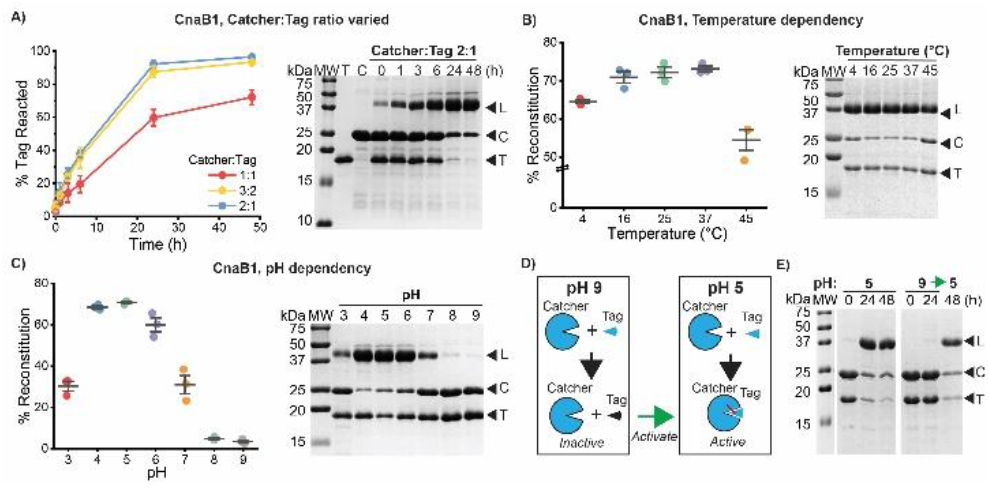
Optimization of CnaB1Catcher/Tag(S6/7a) ligation. A) Adding excess of one of the precursors can drive reaction to completion. B) and C) Temperature and pH dependency of the reaction. D) and E) Schematic presentation and SDS-PAGE analysis showing recovery of the ligation reaction upon changing pH. M stands for molecular weight marker. L, C, and T stand for ligation product (H6-Smt3-Catcher-Tag-CBM-H6), Catcher-precursor (H6-Smt3-Catcher) and Tag-precursor (Tag-CBM-H6), respectively.

To study ligation activity of the newly designed pair *in vivo*, the two precursors coding for the Catcher and Tag were subcloned into vectors carrying different origin of replications and antibiotic resistances for coexpression ^[36]^. (Figure 4A). Ligation product was observed despite the pH ∼7.4-7.8 maintained in *E. coli* cytoplasm ^[37]^ being non-optimal (Figure 4B). Adjusting the expression levels of the two precursors to overexpress the Tag-precursor without an H6-tag drove the ligation reaction of the Catcher-precursor into near completion allowing us to purify only the ligation product (Figure 4B).

**Figure 4.**
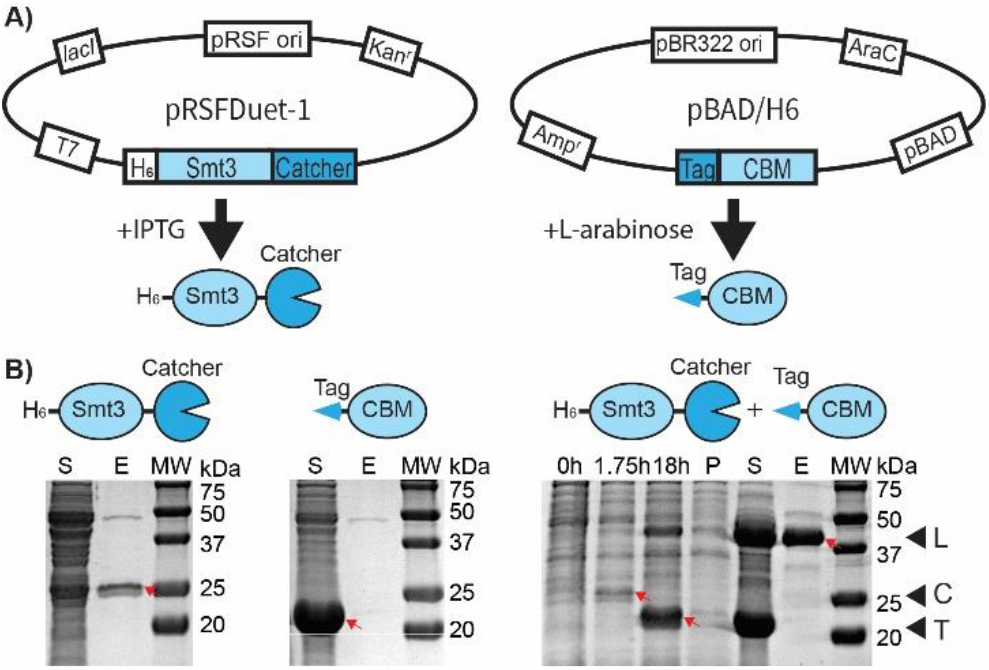
*In vivo* ligation with CnaB1Catcher/Tag(S6/7a). A) Schematic presentation of the plasmid construction. B) SDS-PAGE gels showing expression of the two precursors alone and as coexpression. Timescale of the expression is shown for the coexpression. SilkTag-CBM was induced by adding arabinose at 0h, H6-Smt3-SilkCatcher was induced by adding IPTG at 1.75h. S stands for supernatant, E for elution, MW for molecular weight marker, and P for pellet. L, C, and T, stand for ligation product, Catcher, and Tag which also marked as red arrows.

### Analysis of split variants

We wanted to explore how the split site affects ligation yield and compared the capabilities of four Catcher/Tag variants, split at four different loops, to form an isopeptide bond (Figure 5). Similar to the naming of the original split sites, we named the split sites according to the β-strands flanking the loop that was split, and call them sites S1/2, S4/5, S5/6, and S6/7. The longer one of the two split fragments is always called Catcher and the shorter one Tag. Catcher/Tag pairs are typically engineered by splitting the CnaB domain either at the loop corresponding to that of the split site S1/2 (SnoopCatcher/Tag, ^[15]^) or that of split site 6/7 (SpyCatcherTag, ^[6]^) in Figure 5A. We systematically tested splitting at four different loops (Figure 5A-C). In addition, we tested variation of the original S6/7a split site, named S6/7b, which was split 5 residues closer to the C-terminus. In addition, S6/7b Tag (N6/7b) contains 3 residues less from the C-terminal side. While minor ligation activity was observed for all the split variants, only the two Catcher/Tag pairs split at the last loop (S6/7) were found to have high yields >60% (Figure 5C-E). The yield from the S6/7b split variant was significantly worse than that of the S6/7a (Figure 5D), despite the difference in split site is just 5aa. Interestingly, mixing the S4/5 Tag, which was not able to react with the S5/6 Catcher, with either of the Catchers split at the S6/7 site, improved the ligation yield (Figure 5D). Studying cross-reactivity of the different split variants showed that whereas only short gaps can be tolerated, major overlap of the fragments does not prevent the reaction (Figure 5E). The Catcher/Tag pairs split at the S6/7 loop did cross-react with each other and the 3 residues missing did not prevent the reaction. However, any larger gap between the Catcher and Tag fully abolished the reaction. Instead, major overlap of the Catcher and Tag fragments did not prevent the ligation reaction. The two Catchers split at the S6/7 site were able to mediate ligation with all the tested Tags. Even the S1/2 Catcher and the S6/7b Catcher, which have a 71aa overlap, retained ligation activity together. Similar results were reported for the *E. coli* glycinamide ribonucleotide formyltransferase and, as an extreme example, in the domain swapping of inteins ^[38,39]^.

**Figure 5.**
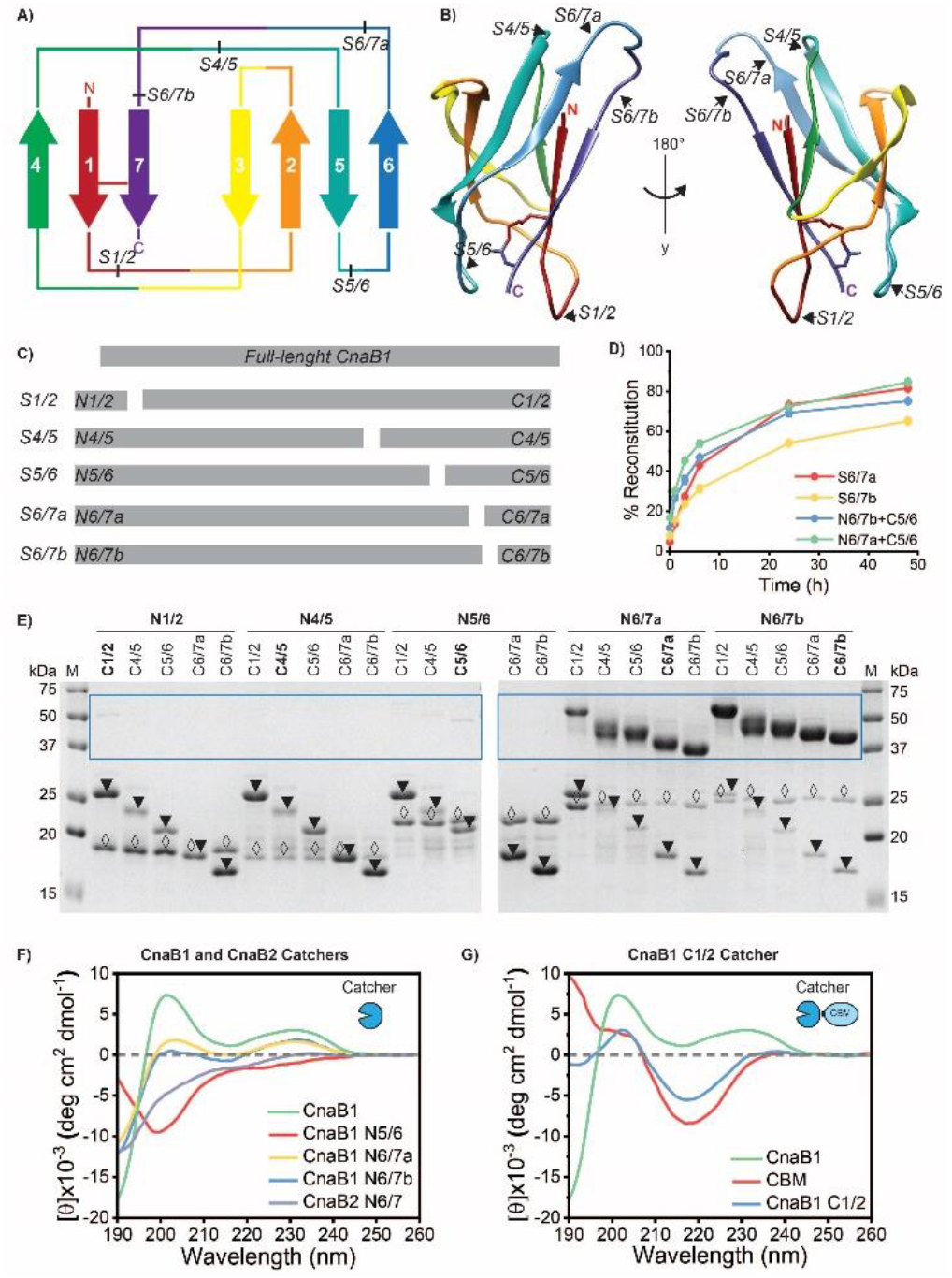
Split sites in CnaB1. A) and B) Different split sites shown in the topology map (A) and cartoon presentation (B) of the homology model of CnaB1. C) Schematic presentation of the lengths and the naming of the split fragments. D) Plot showing the reaction kinetics of the S6/7a, S6/7b pairs, and the N6/7a and N6/7b with C5/6. E) Results from cross-reactivity tests of the different split fragments. M stands for molecular weight marker. The area in which the ligation products appear is highlighted with a blue box, N- and C-precursor are depicted with open diamonds and filled triangles, respectively. F) and G) CD spectra of selected CnaB1 and CnaB2 Catchers. Full-length CnaB1 is shown as a reference.

To better understand the differences in ligation activities, we studied the folding of the Catcher variants using Far-UV CD spectroscopy. CnaB domain has a characteristic CD spectrum with two positive peaks at ∼200 nm and at ∼230 nm (Figure 5F). The CD spectra of the two functional CnaB1 Catchers split at the last loop (S6/7) indicated that they were partially folded, showing smaller positive peaks at both ∼200 nm and ∼230 nm (Figure 5F). Instead, significantly less folded structures were observed for the Catcher variants split at the S5/6 and at the S1/2 loops (Figure 5F, G). The Catcher variant split at the S4/5 loop could not be analyzed due to poor solubility after cleaving the solubility tag. In addition, the CnaB2 Catcher split at the same S6/7 loop was mainly random coil (Figure 5F). Splitting deeper to the protein often promotes association but may also lead to misfolding or prevent folding because of the hydrophobic regions exposed ^[34]^. We interpret that despite having some degree of structure, the two Catchers are not in a fully folded state, because significant increase in the intensity of the two positive peaks is observed to take place along the ligation reaction (Figure 2C). The data indicates that partial folding, perhaps into a molten globule type structure, improves protein fragment complementation and thus also ligation yield. It is known that sites essential for folding are typically not suitable for splitting ^[34]^, which can explain why no ligation was observed when split at the loop closest to the N-terminus (S1/2). In addition to lower ligation yields, the Catchers showing less folded structures (S1/2, S4/5 and S5/6 as well as CnaB2(S6/7) Catcher) were found to be less soluble (data not shown). The solubility of the split fragments is an important factor for biotechnological applications in addition to affecting the ligation yields.

### Ligation of spider silk proteins

Large-scale production of spider silk proteins has been hindered by difficulties in the recombinant production of native-sized spider silk proteins in high yields and in a soluble form ^[21]^. Attempts to express long dragline silk sequences in production organisms have generally resulted in truncated proteins, low expression yields, and/or insoluble proteins although the fibers spun from them have had promising properties ^[24,40,41]^. Therefore, we employed an alternative strategy, in which the shorter but soluble spidroins are covalently conjugated *in vitro* using isopeptide bond formation as initially demonstrated with the SpyCatcher/Tag protein-peptide pair. We have previously shown that SpyCatcher/Tag could be used to ligate 2-3 of 43 kDa fragments of the dragline silk repeat sequence ADF3 from *Araneus diadematus* (ADF3) ^[42,43]^. We next wanted to test, if we could apply the newly engineered Catcher/Tag pair (CnaB1 S6/7) for the ligation of longer spider silk proteins. We decided to call this SilkCatcher/Tag pair. Two prerequisites for the success of the strategy are that the SilkCatcher/Tag pair should not be cross-reactive with the SpyCatcher/Tag-pair and it should be possible to produce in high yields and in a soluble form as a fusion protein with silk. We first studied cross-reactivity with model proteins and no cross-reactivity between the SpyCather/Tag and SilkCather/Tag pairs was observed (Figure S4). Next, we constructed fusion proteins of the ADF3 repeat sequence ^[43]^ with the SilkCather and a cellulose binding module (CBM) from the *Clostridium thermocellum* cellulosome ^[44]^. Fusion with CBM improves solubility of the silk protein and allows utilizing it as a composite material with cellulose as well as an adhesive on delignified cellulose ^[45,46]^. The fusion protein could be expressed in high yields and purified by heat precipitation at 70°C (Figure S5). Attempt to use SnoopCatcher as a reference showed that the combination with ADF3 was not soluble and was therefore not pursued further.

We designed two sets of three precursors (Figure 6A) that allowed us to covalently ligate 4× and 5× silk proteins using a combination of the SpyCatcher/Tag and SilkCatcher/Tag pairs. Five-fragment conjugation could be achieved with only three precursors because Catcher and Tag can be fused in both the N- and C-termini of the target protein ^[25]^. The pH optimum of the SilkCatcher is very close to that of SpyCatcher, making it easy to find suitable conditions to conduct one-pot ligation (Figure 3 and ^[11]^). Indeed, we were able to ligate 4× and 5× silk proteins in a 48h one-pot ligation (Figure 6B, C). The yield from especially the 5× silk protein ligation reaction was, however, not satisfying. Attempts to optimize the ligation conditions further did not improve the yield. We have observed that the ligation efficiency of Catcher/Tag pairs may vary dependening on the other parts of the fusion proteins. Fusion proteins with silk were more prone for aggregation than those with the model proteins and thus more sensitive to the ligation conditions. This might explain why the obtained yield was lower than theoretically expected.

**Figure 6.**
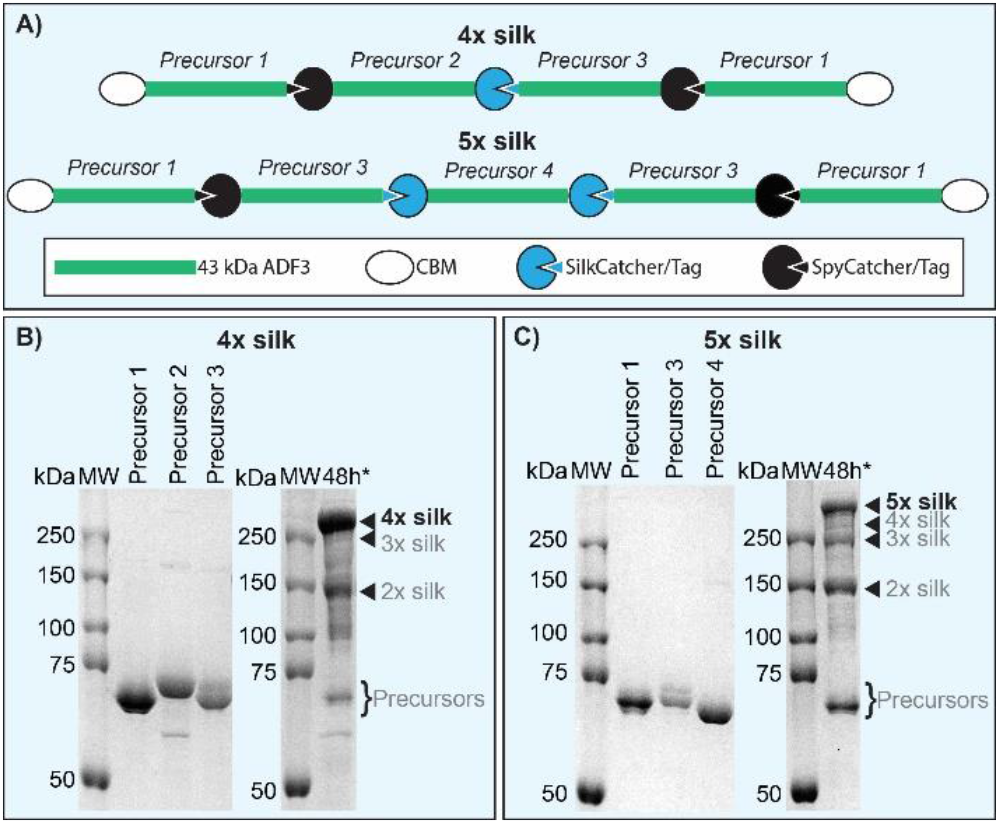
*In vitro* ligation of 4× and 5× silk. A) Schematic presentation of the constructs and the ligation strategy. B) SDS-PAGE gels showing ligation of 4× and 5× silks.

In order to overcome the limitations with yield and to minimize the required purification steps, we decided to take an alternative approach. The approach is based on the following three concepts: 1) the possibility to drive the reaction yield close to 100% by adjusting the reaction ratios (Figure 3A), 2) observation that SilkCatcher/Tag was active in cell lysate, and 3) the design of a set of constructs with and without H6-tags (Figure 7). Ligation is performed in two steps, first in cell lysate with the SilkCatcher/Tag pair, followed by IMAC purification and subsequent ligation using the SpyCatcher2/Tag pair (Figure 7A). In step 1, all precursors 1-4 were expressed in *E. coli*. After harvesting the cells, cell pellets of the H6-tagged precursor 2 and non-tagged precursors 1 and 3 were mixed in 1:2 ration, respectively, so that the precursors with H6-tags were consumed in the ligation reaction. Cell pellet mixtures were lysed and incubated at 37 °C for 48h prior to purification of the ligation product by IMAC. Because almost no precursors with H6-tag were left, only the ligation product was obtained in the elution fractions (Figure 7B). In the second step, ligation product from step 1 were mixed *in vitro* either with each other or with precursor 4 to result in 4× and 5× silks, respectively. Careful adjustment of the precursor ratios and ligation conditions resulted in the reaction to proceed into near completion during 1h incubation with 93.1% ±1.1%, and 92.5%±0.9% ligation yield (Figure 7C).

**Figure 7.**
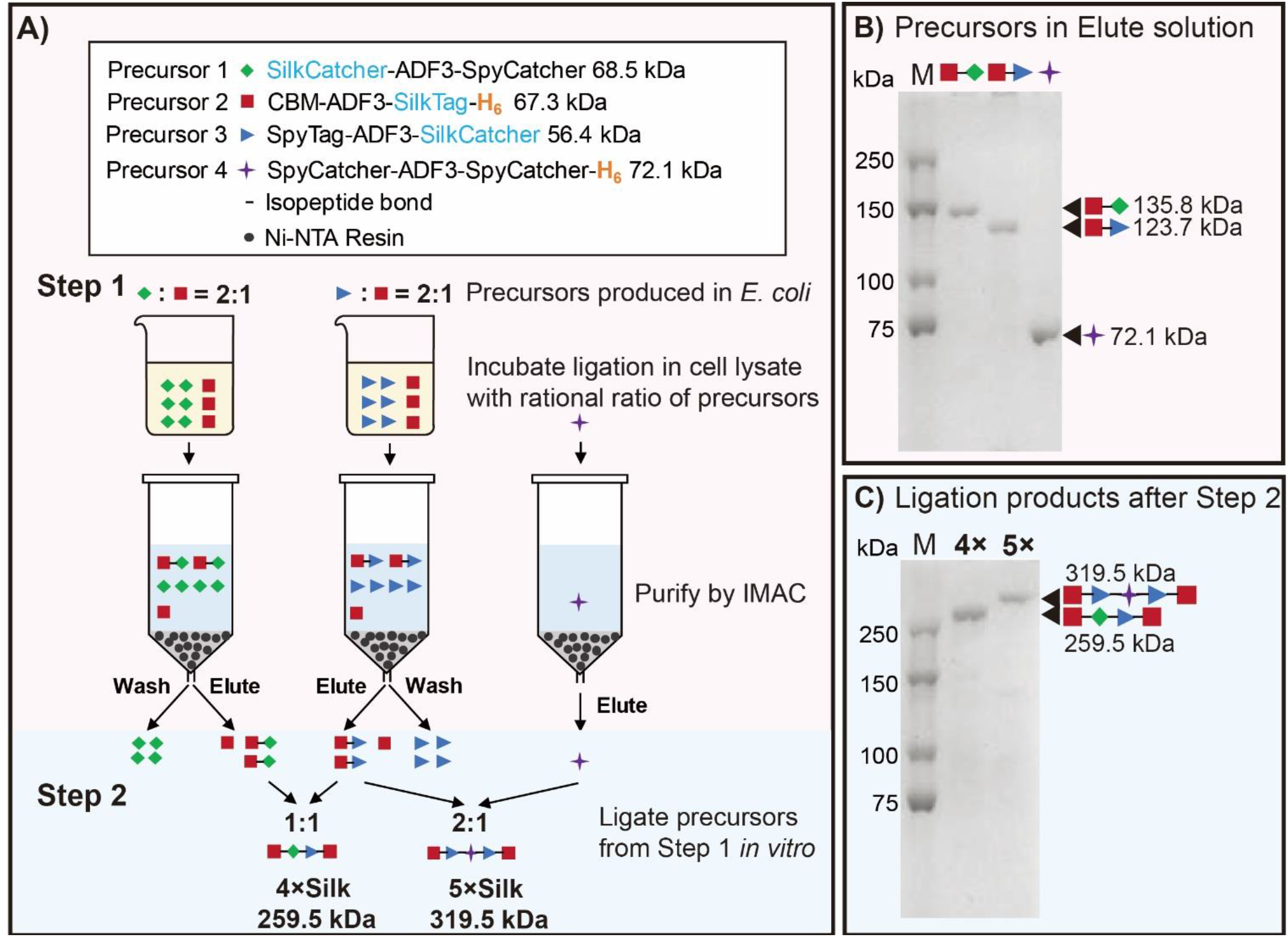
Stepwise ligation of 4× and 5× silk. A) Schematic presentation of the constructs and ligation strategy for 4× and 5× silk. B) An SDS-PAGE gel showing the IMAC purified ligation products after step 1. C) SDS-PAGE gel showing ligation products of step 2 after 1h incubation. M stands for protein weight marker.

## Conclusion

We reported here the design of novel Catcher/Tag pairs starting from two CnaB domains of a cell surface protein of a gut bacterium. We solved the crystal structure of one of the CnaB domains in 1.86 Å resolution which confirmed the existence of an isopeptide bond and assisted the design of the split variants.

The most promising of the pairs was used to covalently ligate native-sized silk proteins consisting of 4 and 5 of 43 kDa fragments of spider silk repeat sequence. Although the yield from the one-pot *in vitro* reaction was only moderate, due to the many ligation reactions and the compromised efficiency of the newly engineered SilkCatcher/Tag pair, we were able to achieve >90% ligation yield by applying a sequential ligation approach. The conjugation of spidroins using the SpyCatcher/Tag protein peptide-pair has significant advantages over the previously reported conjugation through dimerization *via* disulfide bridges or by the protein *trans*-splicing of inteins ^[41,47–51]^. Both the disulfide-bridge and intein-mediated ligation are dependent on the redox state of the system, which limits their applications. In addition, all the reported cases of production of the >50 kDa spidroins were performed using eggcase, flagelliform, or aciniform silks, which are more soluble but do not have comparable mechanical properties to dragline silks. Further optimization of the SilkCatcher/Tag pair may further widen the range of possible applications. The newly designed SilkCatcher/Tag pair is efficient, robust, and orthogonal with the widely used SpyCatcher/Tag pair, and thus a valuable addition to the toolbox of Catcher/Tag pairs.

## Experimental Section

Please see the Supplementary Information for details.

## Acknowledgements

We thank Eva Crosas for her help with the early phases of the structure refinement. Protein crystallization was performed at SPC facility at EMBL Hamburg and the CD spectroscopy and mass spectrometry of the crystallized proteins at the Center for Structural Systems Biology (CSSB, Deutsches Elektronen-Synchrotron DESY). We acknowledge technical support by the SPC facility at EMBL Hamburg. The synchrotron data was collected at beamline operated by EMBL Hamburg at the PETRA III storage ring (DESY, Hamburg, Germany). This work was supported by the Academy of Finland through its Centres of Excellence Programme Life-Inspired Hybrid Materials (LIBER, 2022–2029) under project no 346105 and Academy of Finland projects nos. 317395, 308772, and 333238. We are grateful for the support by the FinnCERES Materials Bioeconomy Ecosystem and use of the Bioeconomy Infrastructure at the Aalto University.

## Entry for the Table of Contents

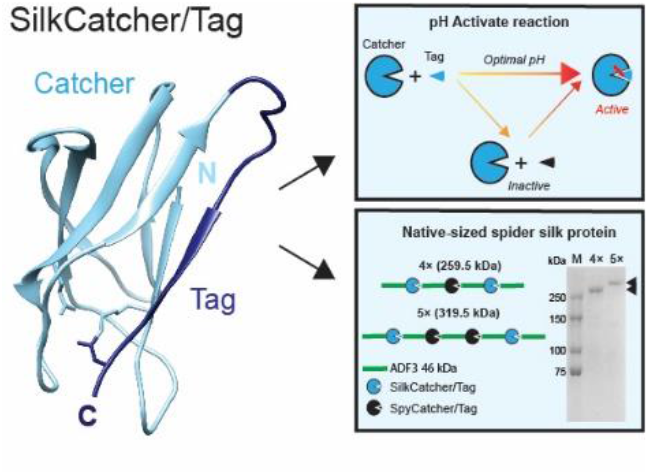

A novel Catcher/Tag pair to be used for isopeptide-bond mediated biological Click-reactions. The newly designed pair, named SilkCatcher/Tag pair, shows activity both *in vivo* and *in vitro* and is pH switchable. Native-sized spider silk-like protein in very high purity is produced here by combining SilkCatcher/Tag pair with the widely used SpyCatcher/Tag pair.

