## supplemental file for "Engineering a minimal SilkCatcher/Tag pair compatible with SpyCatcher/Tag pair for the production of native-sized spider silk"

**Abstract:** Protein/peptide pairs, called Catcher/Tag pairs, are applied for biological isopeptide-bond mediated Click-reactions. Covalent protein ligation using Catcher/Tag pairs has turned out to be a valuable tool in biotechnology and biomedicines. It is essential to increase the current toolbox of Catcher/Tag pairs to expand the range of applications further, e.g., for multiple-fragment ligation that requires several orthogonal ligases. We report here engineering of novel Catcher/Tag pairs for protein ligation, aided by a new crystal structure of a minimal CnaB domain from *Lactobacillus plantarum*. We engineer several split variants, characterize in detail one of them, named SilkCatcher/Tag pair, and show that the newly engineered SilkCatcher/Tag pair is orthogonal and compatible with the widely used SpyCatcher/Tag pair. Finally, we demonstrate the use of the new SilkCatcher/Tag pair in the production of native-sized highly repetitive spider-silk-like proteins with >90% purity, which is not possible with the traditional recombinant production.

### Experimental Procedures

#### Molecular Cloning

Molecular cloning of each is described below and primer sequences are shown in Table S1.

CnaB1 and CnaB2 domains were amplified from a synthetic gene (GeneArt) codon optimized for *E. coli* using primers #SA142 and #SA145 for CnaB1 and #SA143 and #SA144 for CnaB2. The PCR products were digested with *Bam*HI and *Xho*I and inserted into a pEt28-vector coding for N-terminal fusion of H6-Smt3[1], resulting in plasmids pSAEt190 (CnaB1) and pSAEt191 (CnaB2).

SilkCatcher and SilkTag were amplified from pSAEt190 using primers #SA161 and #SA163 (Catcher) and #SA160 and #SA162 (Tag). SilkCatcher was digested with *Bam*HI and *Xho*I and inserted into H6-Smt3 plasmid[1] to result in pSAEt200. SilkTag was digested using *Nco*I and *Eco*RI and inserted into pSAEt42[2] in plasmid pSAEt201. Genes coding for LplCnaB2Catcher(S6/7), LplCnaB2Tag(S6/7), and LplCnaB2Tag(S6/7)_D91A were ordered from GeneArt and inserted into plasmids H6-Smt3[1] (Catcher, *Bam*HI and *Xho*I) and pSAEt42 [2] (Tags, *Nco*I and *Eco*RI) to result in plasmids pSAEt231, pSAEt241, and pSAEt232, respectively. Constructs pSAEt238(CnaB1Catcher(S6/7b)), pSAEt239(CnaB1Tag(S6/7b)), pSAEt214(CnaB1Catcher(S5/6)), and pSAEt215(CnaB1Catcher(S1/2)) were constructed by ordering genes coding for the respective CnaB1Catchers and CnaB1Tag from GeneArt and inserting them into the vectors H6-Smt3[1] (Catchers, *Bam*HI and *Xho*I) and pSAEt42[2] (Tag, *Nco*I and *Eco*RI), respectively. To construct pSAEt217 and pSAEt213, the longer CnaB1Tags were amplified from the synthetic gene coding for CnaB1 using primers #SA161 and #SA173 (pSAEt217) or #SA161 and #SA171 (pSAEt213) and inserted into H6-Smt3 between *Bam*HI and *Xho*I sites. Plasmid pSAEt216 was constructed by amplifying CnaB1Tag(S4/5) using primers #SA172 and #SA160 from the synthetic gene and inserting it into vector pSAEt42[2] between *Nco*I and *Eco*RI sites.

pSALBAD306 was constructed by digesting pSAEt200 using restriction enzymes *Nco*I and *Xho*I and ligating the insert into plasmid pSABAD289. Plasmid pSABAD289 was modified from pBAD/HisA (GeneArt) in order to introduce *Xho*I site in front of *Hind*III site. pSARSFDuet307 was constructed by transferring the sequence coding for H6-Smt3-SilkCatcher from pSAEt201 into pRSFDuet-1 vector (Novagen) between *Nco*I and *Xho*I sites.

To construct pSAEt338 and pSAEt346, genes coding for CnaB1Tag(S6/7a) carrying the D91A or D92A mutation, respectively, were ordered from GeneArt and inserted into vector pSAEt42 [2] between *Nco*I and *Eco*RI sites.

**Table S1.** List of primers.

| Primer name | Primer sequence |
|---|---|
| #SA27 | 5'-ACAGAATTCTAGCTCCGCACATATTGTT |
| #SA77 | 5' -CGCCATGGGTTCCCTTGTTCCTGAATTA |
| #SA78 | 5' -CGCTCGAGTTTTAAAGCGTCGGTTAAAA |
| #SA142 | 5'-CCGGATCCATGACCTATACCATTGAAC |
| #SA143 | 5'-CCGGATCCACCGGTGATGTTACCCTGAC |
| #SA144 | 5'-GTGCTCGAGCTATCCCGGTTCGGTAACC |
| #SA145 | 5'-GTGCTCGAGCTAGGTCGGAACAACCGGC |
| #SA160 | 5'-CTAGAATTCGCGGTCGGAACAACCG |
| #SA161 | 5'-CCGGATCCACCTATACCATTGAACTG |
| #SA162 | 5'-ATACCATGGGCATTAAACCGGAAGTTGC |
| #SA163 | 5'-GTGCTCGAGCTAACCTAAGGTAAATTTC |
| #SA171 | 5'-GTGCTCGAGCTAGGCACGCAGATTTTTAAC |
| #SA172 | 5'-ATACCATGGGTGATTATCAGTTTGTGG |
| #SA173 | 5'- GTGCTCGAGCTACAGCGGTGCTTTTG |
| #SA174 | 5'-ATACCATGGGTTATGAGCTGAATACC |
| #SA183 | 5'-GGAGATATACCATGGGCACCTATA |
| #SA184 | 5'-GCGCTAGCGCTCGGGGATCCTAAGGTA |
| #SA185 | 5'-GCGAATTCTAGCTCCGGCATTAAACCG |
| #SA191 | 5'-GTGCTCGAGACCGGTACCGGTCGGAACAACC |
| #SA204 | 5'-CCAGACTCGAGTTATTTGGTCGG |

Plasmid pSARSFDuet311 was constructed stepwise. First, CBM between sites NcoI and NheI in plasmid pSAEt56[2] was replaced with CnaB1Catcher(SilkCatcher, S6/7a) by amplifying CnaB1Catcher(SilkCatcher, S6/7a) from plasmid pSAEt190 using primers #SA183 and #SA184, to result in pSAEt266. Fragment between *Nco*I and *Xho*I was then transferred into vector into pRSFDuet-1 vector (Novagen), to result in pSALRSFDuet310. Finally, pSALRSFDuet311 was constructed to introduce a stop codon after SpyTag by amplifying SpyTag from a pSAEt56 using primers #SA027 and #SA204. The PCR product was digested by restriction enzymes *Eco*RI and *Xho*I and ligating the insert into plasmid pSALRSFDuet310, to end up in pSALRSFDuet311 coding for CnaB1Catcher(SilkCatcher, S6/7a)-ADF3-SpyTag.

pSAEt263 coding for CBM-ADF3-CnaB1Tag(SilkTag, S6/7a)-H6 was constructed by amplifying CnaB1Tag(SilkTag, S6/7a) from plasmid pSAEt190 using primers #SA185 and #SA191, followed by digestion with *Eco*RI and *Xho*I and ligation into vector pSAEt42[2]. pSAEt276 was constructed by stepwise replacement of the two CBMs in pSAEt42 with CnaB1 Tag(S6/7a) Catcher and Tag. First, CnaB1Catcher(S6/7a) was amplified with primers #SA183 and #SA184 from plasmid pSAEt190 and inserted into plasmid pSAEt42 between *Nco*I and *Nhe*I sites, followed by replacing the C-terminal CBM with a synthetic gene (GeneArt) coding for CnaB1 Tag(S6/7a) digested with *Eco*RI and *Xho*I to result in plasmid coding for CnaB1Catcher(SilkCatcher, (S6/7a))-ADF3- CnaB1Catcher(SilkCatcher, (S6/7a)).

pSALRSFDuet300 was constructed by replacing the C-terminal SpyCatcher in pYY3c with CnaB1(SilkTag, S6/7a)Tag-H6. A gene coding for the Tag-H6 was ordered from GeneArt and subcloned into pYY3c between *Eco*RI and *Xho*I sites to results in SpyCatcher2-ADF3-CnaB1(SilkTag, S6/7a)-H6. pSALRSFDuet317 was constructed by digesting pSAEt253 using restriction enzymes *Eco*RI and *Xho*I and ligating the insert into plasmid pSALRSFDuet300.

Gene coding for Ulp1 was amplified from Saccharomyces cerevisiae S3328 cells with primers #SA77 and #SA78, followed by insertion into vector pSAEt28A between *Nco*I and *Xho*I sites, to result in plasmid pSAEt128 coding for Ulp1-H6. Construction of plasmids pSAEt56[2]and pYY3c[3] has been reported before.

#### Protein expression and purification (Ulp1)

*E. coli* T7 express strain (NEB) was transformed with plasmid pSAEt128 coding for Ulp1-H6. Cells were grown in 0.3 L autoinduction media (Magic Media, ThermoFisher) supplemented with kanamycin (50 µg/mL final concentration) according to the manufacturer's protocol and harvested by centrifuging 4,670 ×g for 10 mins at 15°C. Cell pellets were resuspended into Buffer A (50 mM NaPi pH 8.0, 300 mM NaCl) and lysed using Emulsiflex C3 (Avestin). The soluble fraction was collected by centrifuging 15,000 ×g for 15 mins and purified by IMAC using 5 mL HisTrap FF column (Cytiva). The elution fractions containing the desired protein were pooled and buffer exchanged into 25 mM Tris-HCl pH 7.4 using Econo-Pac 10DG columns (BioRad).

#### Protein expression and purification (full length CnaB1 and CnaB2)

*E. coli* T7 express strain (NEB) was transformed with plasmids pSAEt190 and pSAEt191 coding for H6-Smt3-CnaB1 and H6-Smt3-CnaB2, respectively. Cells were grown in 0.2L autoinduction media (Magic Media, ThermoFisher) supplemented with kanamycin (50 µg/mL final concentration) according to the manufacturer's protocol and harvested by centrifuging 4,670 ×g for 10 mins at 15°C. Cell pellets were resuspended into Buffer A (50 mM NaPi pH 8.0, 300 mM NaCl) and lysed using Emulsiflex C3 (Avestin). The soluble fraction was collected by centrifuging 15,000 ×g for 15 mins and purified by IMAC using 5mL HisTrap FF column (Cytiva). The elution fractions containing the desired protein were pooled and buffer exchanged into MQ using Econo-Pac 10DG columns (BioRad). The H6-Smt3 solubility tag was removed by digestion with Ulp1 (homemade). The digested protein was rerun into a 5 mL HisTrap FF column and the flow through fractions containing the CnaB domains were collected. Following a buffer exchange into MQ using Econo-Pac 10DG columns (BioRad) columns the proteins were concentrated using Vivaspin centrifugal concentrators (Sartorius) with cutoff of 5,000 MWCO.

#### Protein expression and purification (split CnaB1 and CnaB2)

*E. coli* T7 express strain (NEB) was transformed with plasmids coding for the Catcher/Tag fusion proteins. Cells were grown in 0.2 L autoinduction media (Magic Media, ThermoFisher) supplemented with kanamycin (50 µg/mL final concentration) according to the manufacturer's protocol and harvested by centrifuging 10,000 ×g for 10 mins at 4 °C. Cell pellets were resuspended into Buffer A (5 mM NaPi pH 8.0, 300 mM NaCl) and lysed using sonicating (Q500 Sonicator, Qsonica). The soluble fraction was collected by centrifuging 20,000 ×g for 15 mins at 4 °C and purified by IMAC using 5 mL HisTrap FF column (Cytiva). The elution fractions containing the desired protein were pooled and buffer exchanged into MQ using Econo-Pac 10DG columns (BioRad). Finally, the proteins were concentrated using Vivaspin centrifugal concentrators (Sartorius) with cutoff of 5,000 MWCO.

#### Protein crystallization

Crystals of CnaB2 were obtained using protein solution with concentration of 46 mg/ml in MQ water. Diffraction data was collected from crystals prepared manually by mixing protein and reservoir solution 1:1 in a 4 ul drop and equilibrating against 500 ul of reservoir solution in a hanging-drop vapour-diffusion experiment. The best diffracting crystals were obtained from 1.5 M ammonium sulfate, 0.1 M Tris-HCl pH 8.5 at RT. Crystals were cryo-protected by quickly soaking into reservoir solution containing 30 % of glycerol prior to flash-cooling with liquid nitrogen.

#### Data processing and structure refinement

Diffraction data for the CnaB2 crystals were collected on P13 beamline at EMBL Hamburg, PETRA III[4]. The data were processed with *XDS*[5] and scaled with *Aimless*[6–8], from the CCP4 software suite[9], to 1.86 Å resolution. Due to the packing of the molecules into a crystal packing forming a hollow sphere, only 12 molecules were packed in the ASU instead of >20 predicted based on the solvent content. The estimated Matthews coefficient $V_M$ [10] for the 12 molecules in the ASU was 4.00 Å$^3$/Da, corresponding to 69.3 % solvent content $V_S$. The initial structure was solved with molecular replacement by a homology model constructed with *SwissModel* [11] using PDB 3PHS [12] as a starting model. The solution could be obtained after trimming the loops of the homology model and constructing a trimeric model based on the initial searches using *Phaser* software[13]. By searching for all possible space groups in the same pointgroup the space group was defined to be $P3_221$ instead of the $P3_121$ assigned initially by pointless [6,7], after which *FREEFLAG* [14,15] was used for calculating 5 % of free R factors for the data set. *ARP/wARP* [16] was used to build the missing fragments to the initial solution from *Phaser*, followed by several rounds of refinement using Refmac5[17]. Further refinement was done with *phenix.refine* [18] from the *PHENIX* software suite [19] in iterative cycles of manual model building in *COOT* [20]. Isotropic individual temperature factors were refined. In the final stages of the refinement, 67 TLS groups were defined by *phenix.find_tls_groups* tool [18] and added to the refinement parameters. Data collection, processing and structure refinement statistics are listed in Table S2.

**Table S2.** Data collection and structure refinement statistics of *L. plantarum* CnaB2 domain

| | CnaB2 |
|---|---|
| Beamline | EMBL P13 |
| Wavelength (Å) | 0.9802 |
| Space group | $P3_221$ |
| Molecules/ASU | 12 |
| Unit cell *a*, *b*, *c* (Å); | 120.06, 120.06, 230.92 |
| $\alpha$, $\beta$, $\gamma$ (°) | 90, 90, 120 |
| Resolution (Å) | 104-1.86 (1.93-1.86) |
| $R_{merge}$ (%) | 7.68 (84.8) |
| CC 1/2 | 99.9 (73.0) |
| No. of unique reflections measured | 158616 (13576) |
| Mean $I/\sigma(I)$ | 17.85 (1.18) |
| Wilson B-factor | 30.34 |
| Completeness (%) | 98.46 (85.01) |
| Multiplicity | 9.4 (4.5) |
| **Refinement** | |
| Resolution (Å) | 104-1.86 |
| No. of reflections refined against | 158501 (13534) |
| $R/R_{free}$ | 0.154/0.187 |
| Mean B (overall Å$^2$) | 36.86 |
| **No. of atoms** | |
| Protein | 8524 |
| Ligands | 214 |
| Water | 1266 |
| **R.m.s. deviations from ideal targets** | |
| Bond lengths (Å) | 0.019 |
| Bond angles (°) | 1.46 |
| **Ramachandran plot (%)** | |
| Favored | 98.84 |
| Allowed | 1.16 |
| Outliers | 0.00 |
| PDB code | 8BDW |

#### *In-vitro* ligation of CnaB1 and CnaB2 Catcher/Tag pairs

Ligation activities of the different CnaB1 and CnaB2 Catcher/Tag variants were performed by mixing 30 µM (final concentration) of each precursor in 1:1 ratio in 50 mM buffer of pH 5.0 (50 mM NaPi pH 7.0 adjusted to pH 5.0 by 50 mM $H_3PO_4$). Reactions were incubated in 37 ℃ for 24 or 48 h. For testing the effect of pH, 50 mM NaPi adjusted to pH 3.0, 4.0, 5.0, 6.0, 7.0, 8.0, or 9.0 were used (50 mM NaPi pH 7.0 adjusted to pH 3.0, 4.0, and 5.0 by 50 mM $H_3PO_4$, 50 mM NaPi pH 7.0 and 8.0, 50 mM NaPi pH 7.0 adjusted to 9.0 by 50 mM $Na_2HPO_4$). For testing the effect of temperature, reactions were incubated in an Eppendorf Thermomixer C at 4, 16, 25, 37, or 45 ℃ for 24 h. For testing the effect of different ratio of precursors, 30 µM H6-CBM-Tag was mixed with 30, 45, and 60 µM H6-Smt3-Catcher, respectively.

All reactions were performed 3 times. All the ligation reactions were quenched by adding SDS-PAGE loading dye and heating at 95℃ for 5 min. The reconstitution reactions were monitored by SDS-PAGE and ligation yields were analyzed by quantifying band intensities from SDS-PAGE gels by Image Lab 6.0.1 software (Bio-Rad). Reconstitution was calculated by dividing the intensity of the band for the ligation product by the sum of intensity of the band for two precursors and ligation product, then multiplying by 100.

#### pH switch

Reactions were performed by mixing each precursor in 30 µM final concentration in 50 mM buffer of pH 5.0 (50 mM NaPi pH 7.0 adjusted to pH 5.0 by 50 mM $H_3PO_4$) and pH 9.0 (50 mM NaPi pH 7.0 adjusted to pH 9.0 by 50 mM $Na_2HPO_4$) and incubated in 37 ℃ for 24 h. Then the pH of the pH 9.0 reaction solution was adjusted to pH 5.0 by 1 M $H_3PO_4$, followed by incubation in 37 ℃ for 24 h more. pH was measured by Micro pH Electrodes (Mettler Toledo).

#### *In-vivo* ligation

Plasmids pSALBAD306 and pSARSFDuet307 were cotransformed into *E. coli* T7 express strain (NEB). Cells were grown in LB-media supplemented with kanamycin (50 µg/mL final concentration) and ampicillin (100 µg/mL final concentration) until OD600 reached 0.4-0.6, at which point expression was induced by adding 0.2% L-arabinose and 0.1 mM IPTG (final concentrations). Cells were incubated at 30°C for 18 h after induction. Samples were collected after 0h, 1.75 h and 18 h by centrifuging the cells at 12,000 ×g for 1 min. After 18 h incubation, cells were harvested and lysed using B-PER Bacterial protein extraction reagent (Thermo Fisher Scientific), followed by purification of the soluble fraction using Ni-NTA spin columns (Qiagen) according to the manufacturer's protocol.

#### CD spectrometry

CD spectra in Figure S1. CnaB1 and CnaB2 domains (from pSAEt190 and pSAEt191 after cleavage of the Smt3) were diluted into 0.27 mg/mL concentration in MQ. Samples were measured using Chirascan CD Spectrometer (Applied Photophysics), for 210-280nm with 1nm bandwith for 0.5s intervals at 22.8℃. Each measurement was repeated three times and the mean value is shown.

Other CD data. To analyze individual proteins, fresh protein samples were diluted into 10 µM concentration in 50 mM buffers of pH 3.0, 5.0, 7.0 and 9.0 (50 mM NaPi pH 7.0 adjusted to pH 3.0 and 5.0 by 50 mM $H_3PO_4$, 50 mM NaPi pH 7.0, 50 mM NaPi pH 7.0 adjusted to pH 9.0 by 50 mM $Na_2HPO_4$) and incubated for 20 min before measurement. To follow the ligation reaction by CD spectrometry, each precursor was mixed in a final concentration of 30 µM in 50 mM buffer pH 5.0 and then incubated at 37 ℃ for 48 h. Samples at different time points were diluted 1:3 with MQ water prior to measurement. All samples were measured using Jasco J-1500 CD spectrometer, for 190-260 nm with 1 nm bandwidth for 0.5 s intervals at room temperature. Each measurement was repeated three times and the mean value is shown.

#### Mass spectrometry

The molecular masses of CnaB1, CnaB2, and the ligation product from the reaction of H6-Smt3-CnaB1Catcher(S6/7a) and CnaB1Tag(S6/7a) were analyzed by mass spectrometry. CnaB1 and CnaB2 were obtained by digesting the fusion protein with Ulp1 as described above. The ligation product was analyzed from the following reaction. The precursors were mixed at 65 µM final concentration in 100 mM NaCl and incubated at RT for 24 h. All three samples were analyzed by MALDI TOF (Bruker/CovalX) at the Center for Structural Systems Biology (CSSB), using sinapic acid as matrix.

#### Protein expression, purification, and *in vitro* one-pot ligation (silk fusion proteins)

Each plasmid coding for the silk fusion-proteins was transformed into *E. coli* T7 express strain (NEB). Cells were grown in 0.5 L EnPresso B500 media (EnPresso) supplemented with kanamycin (50 µg/mL final concentration) according to the manufacturer's protocol and harvested by centrifuging 10,000 ×g for 15 mins at 4 °C. Pellets were resuspended into Buffer A (5 mM NaPi pH 8.0, 300 mM NaCl) and lysed using Emulsiflex C3 (Avestin). The soluble fractions were collected by centrifuging 10,000 ×g for 15 mins at 4 °C. The soluble fractions were purified either by IMAC using 5 mL HisTrap FF column (Cytiva) [CBM-ADF3-SpyTag-$H_6$,

CnaB1Catcher(S6/7a)-ADF3-CnaB1Catcher(S6/7a)-$H_6$, and SpyCatcher2-ADF3-CnaB1Tag(S6/7a)-$H_6$] or by heat fractionation, incubating at 70°C for 20min, centrifuging 10,000 ×g for 15 mins at 4 °C and collecting the soluble fraction [SpyCatcher2-ADF3-CnaB1Catcher(S6/7a)]. The elution fractions containing the desired protein were pooled and buffer exchanged into MQ using Econo-Pac 10DG columns (Bio-Rad). Finally, the proteins were concentrated using Vivaspin centrifugal concentrators (Sartorius) with cutoff of 10,000 MWCO. Concentrations of the precursors were determined based on band intensities from Coomassie blue-stained SDS-PAGE gels using Image Lab 6.0.1 software (Bio-Rad). Concentrations were determined by comparing the intensities to that of an IMAC-purified reference sample of which the concentration had been determined by Nanodrop.

Finally, 4x and 5x silk proteins were ligated by mixing the elution fractions of CBM-ADF3-SpyTag-$H_6$: SpyCatcher2-ADF3-CnaB1Catcher(S6/7a): SpyCatcher2-ADF3-CnaB1Tag(S6/7a)-$H_6$ in ratio of 2:1:1 and CBM-ADF3-SpyTag-$H_6$: SpyCatcher2-ADF3-CnaB1Tag(S6/7a)-$H_6$: CnaB1Catcher(S6/7a)-ADF3-CnaB1Catcher(S6/7a)-H6 in ratio of 2:2:1, respectively. The ligation reactions were performed for 48 h at 37°C with a final concentration of 10 or 20 μM for precursors. All the ligation reactions were quenched by adding SDS-PAGE loading dye and heating at 95°C for 5 min.

**Protein expression, purification, and *in vitro* stepwise ligation (silk fusion proteins)**

Each plasmid coding for the silk fusion-proteins was transformed into *E. coli* T7 express strain (NEB). Cells were grown in 0.5 L EnPresso B500 media (EnPresso) supplemented with kanamycin (50 µg/mL final concentration) according to the manufacturer's protocol and harvested by centrifuging 10,000 ×g for 15 mins at 4 °C. Cell pellets were mixed to achieve 1:2 ratio of precursors [CBM-ADF3-CnaB1Tag(S6/7a)-$H_6$: SpyCatcher-ADF3-CnaB1Catcher(S6/7a) and CBM-ADF3-CnaB1Tag(S6/7a)-$H_6$: SpyTag-ADF3-CnaB1Catcher(S6/7a)]. Concentrations of precursors were estimated based on densitometric analysis of Coomassie blue-stained SDS-PAGE gels using Image Lab 6.0.1 software (Bio-Rad).

Pellets were resuspended into Lysis Buffer (100 mM NaOAc buffer, pH 5.0, 100 mM NaCl, 2 mM EDTA, 1 mM DTT, 1µg/mL DNase I and protease inhibitor) and lysed using Emulsiflex C3 (Avestin). The soluble fractions were collected by centrifuging 10,000 ×g for 15 mins at 4 °C and then incubated at 37 °C for 48 h to ensure complete ligation. The cell pellets of SpyCatcher2-ADF3-SpyCatcher2-$H_6$ were lysed in the same way and the soluble fractions were collected by centrifuging 10,000 ×g for 15 mins at 4 °C. The above ligation products and the soluble fractions of SpyCatcher2-ADF3-SpyCatcher2-$H_6$ were purified by IMAC using 5 mL HisTrap FF column (Cytiva). The elution fractions were pooled, and buffer exchanged into Ligation Buffer (20 mM NaOAc, 20 mM NaCl, pH 5.0) using Econo-Pac 10DG columns (BioRad). After that, the proteins were concentrated using Vivaspin centrifugal concentrators (Sartorius) with cut-off of 10,000 MWCO. Concentrations of the precursors were determined based on band intensities from Coomassie blue-stained SDS-PAGE gels using Image Lab 6.0.1 software (Bio-Red). Concentrations were determined by comparing the intensities to that of an IMAC-purified reference sample of which the concentration had been determined by Nanodrop.

Finally, 4x and 5x silk proteins were ligated by mixing the elution fractions as follows: 1) CBM-ADF3-CnaB1Tag(S6/7a)-$H_6$ and SpyCatcher-ADF3-CnaB1Catcher(S6/7a) in ratio of 1:1, 2) CBM-ADF3-CnaB1Tag(S6/7a)-$H_6$ and SpyTag-ADF3-CnaB1Catcher(S6/7a) in ratio of 1:1, 3) CBM-ADF3-CnaB1Tag(S6/7a)-$H_6$ and SpyTag-ADF3-CnaB1Catcher(S6/7a) in ratio of 2:1, and 4) SpyCatcher2-ADF3-SpyCatcher2-$H_6$ in ratio of 2:1. The ligation reactions were performed for 1 h at 25°C with a final concentration of 10 µM for each precursor. All the ligation reactions were quenched by adding SDS-PAGE loading dye and heating at 95℃ for 5 min. Ligation yields were analyzed by quantifying band intensities by Image Lab 6.0.1 software (Bio-Rad). Reconstitution was calculated by dividing the intensity of the band for the ligation product by the sum of intensity of the band for two precursors and ligation product, then multiplying by 100.

### Results and Discussion

**Figure S1.** CnaB domains in the cell surface protein LP2578 from *Lactobacillus plantarum*.

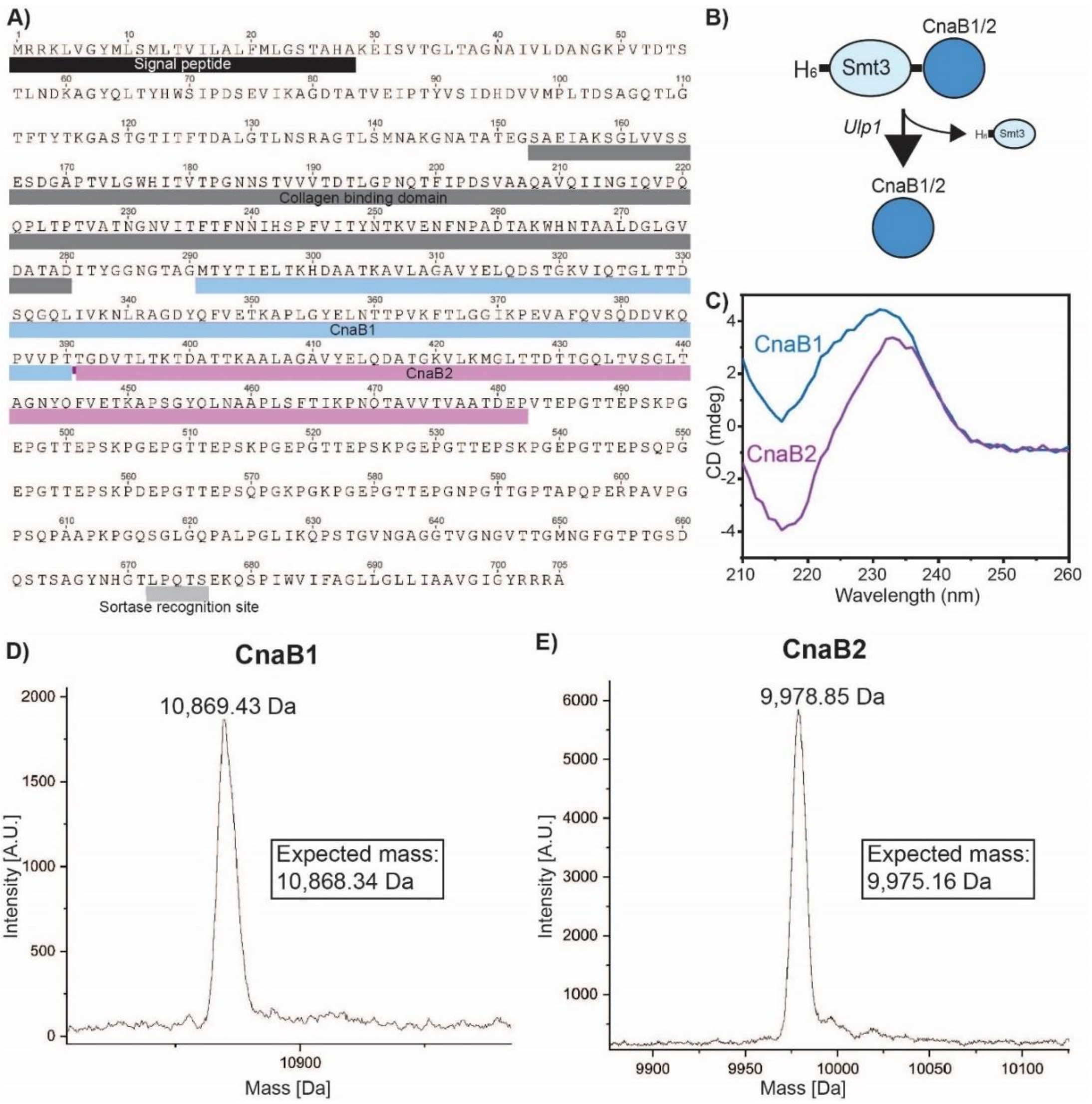

**Figure S1.** CnaB domains in the cell surface protein LP2578 from *Lactobacillus plantarum*. A) The amino acid sequence of LP2578. B) Schematic presentation of the production strategy for CnaB1 and CnaB2. C) CD spectra of CnaB1 and CnaB2. D) and E) Mass spectrometry data indicating the presence of an isopeptide bond in CnaB1 (D) and CnaB2 (E).

**Figure S2.** Comparison of CnaB domains used as starting point for the engineering of Catcher/Tag pairs.

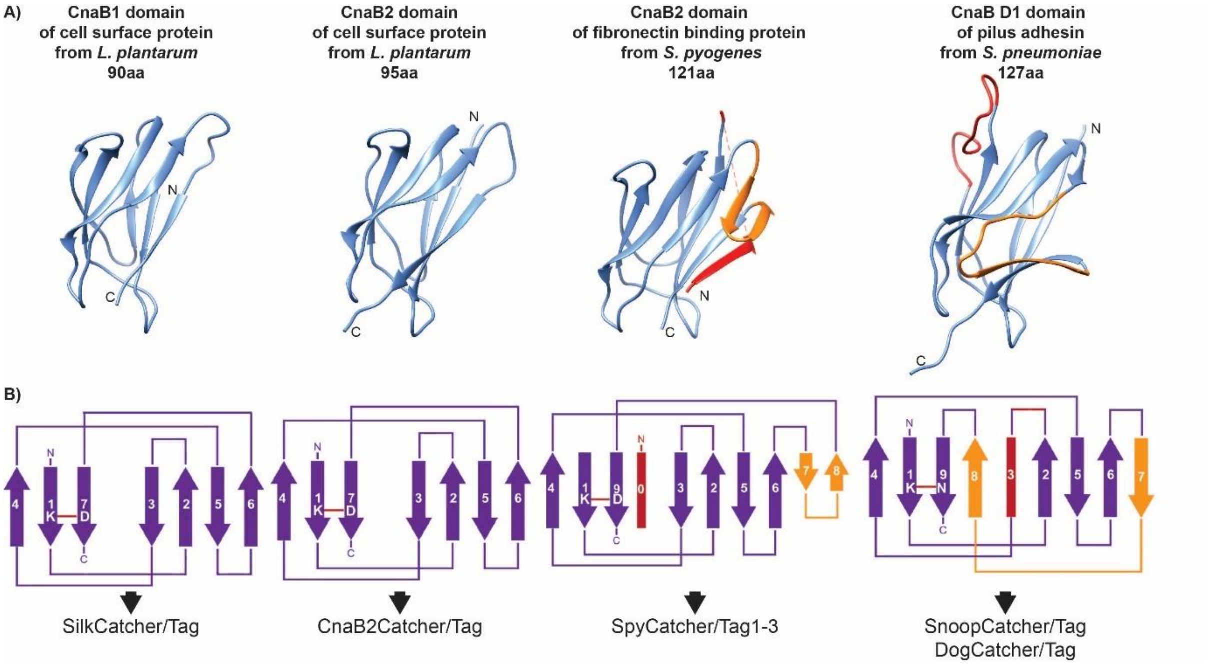

**Figure S2.** Comparison of CnaB domains used as starting point for the engineering of Catcher/Tag pairs. The homology model of CnaB1 from *L. plantarum* and the crystal structures of CnaB2 from *L. plantarum* (this work, PDB ID: 8BDW), *S. pyogenes* CnaB2 domain based (PDB ID: 2X5P), and S. pneumoniae CnaB D1 domain(PDB ID: 2WW8) presented as cartoon models (A) and topology diagrams (B). The regions missing from *L. plantarum* CnaB domains are shown in yellow and red.

**Figure S3.** Ligation reaction of inactive and active CnaB2 Catcher/Tag (S6/7).

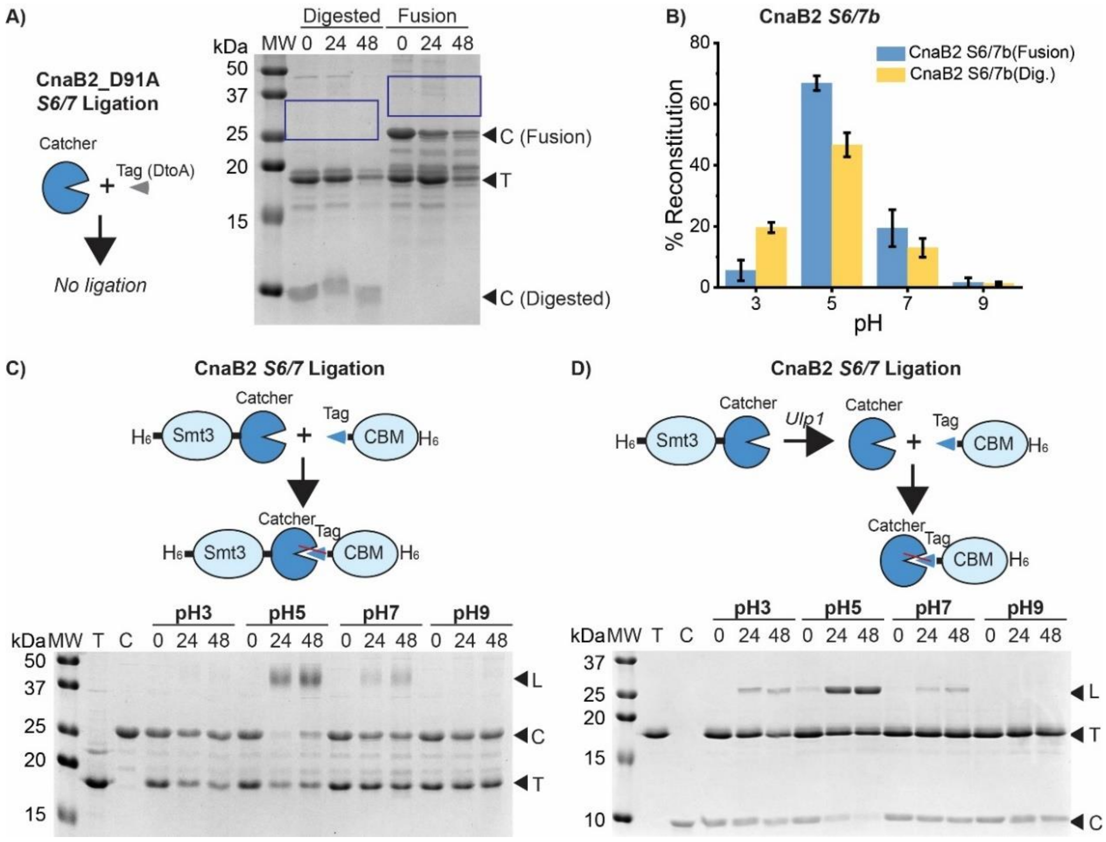

**Figure S3.** Ligation reaction of inactive and active CnaB2 Catcher/Tag (S6/7). A) No ligation product is observed after mixing CnaB2Catcher(S6/7) ("Digested") or H6-Smt3- CnaB2Catcher(S6/7) (Fusion) with CnaB2Tag(S6/7)(D91A)-CBM-H6. B-D) Reconstitution yield (B) and SDS-PAGE gels (C and D) showing reaction of CnaB2Catcher(S6/7) ("Digested") or H6-Smt3-CnaB2Catcher(S6/7) (Fusion) with CnaB2Tag(S6/7)-CBM-H6 at pH 3, 5, 7, and 9. L, C, and T, stand for ligation product, Catcher, and Tag.

**Figure S4.** SilkCatcher/Tag and SpyCatcher2/Tag do not cross-react

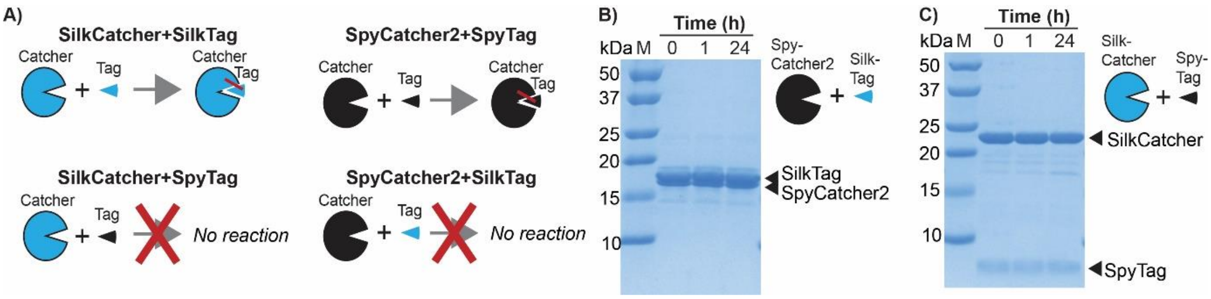

**Figure S4.** SilkCatcher/Tag and SpyCatcher2/Tag do not cross-react. A) Schematic presentation of the B) SDS-PAGE gel showing no ligation takes place upon mixing SpyCatcher2 and SilkTag. C) SDS-PAGE gel showing no ligation takes place upon mixing SilkCatcher and SpyTag.

**Figure S5.** Heat fractionation of a fusion protein of silk and SilkTag or SilkCatcher.

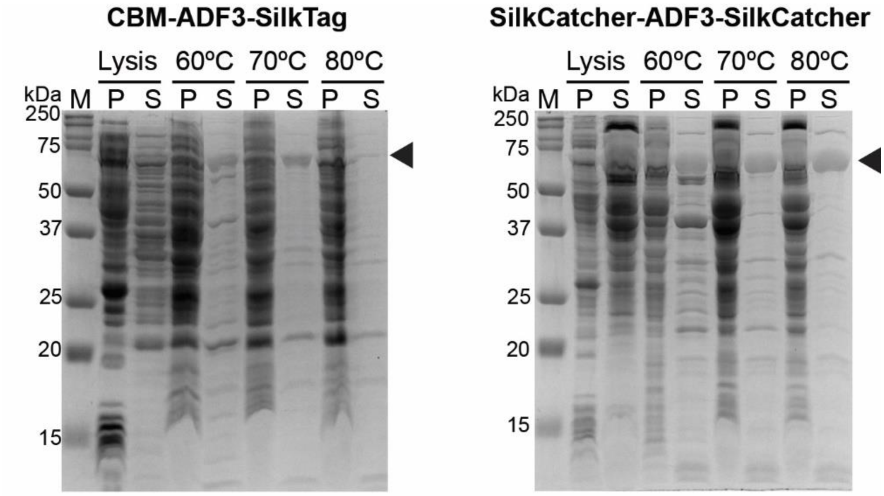

**Figure S5.** Heat fractionation of a fusion protein of silk and SilkTag or SilkCatcher. Cells were lysed, followed by incubating the soluble fraction at 60°C, 70°C, or 80°C for 20min. The fusion proteins remained soluble at 70°C and 80°C, while most *E. coli* proteins precipitated. M, S, and P stand for molecular weight marker, supernatant, and pellet, respectively.
